# Intergenic polyA/T tracts explain the propensity of yeast de novo genes to encode transmembrane domains

**DOI:** 10.1101/2025.03.04.641575

**Authors:** Nikolaos Vakirlis, Timothy Fuqua

## Abstract

New genes can emerge de novo from non-genic genomic regions. In budding yeast, computational predictions have shown that intergenic regions harbor a higher-than-expected propensity to encode transmembrane domains, and this propensity seems to be linked to the high prevalence of predicted transmembrane domains in evolutionarily young genes. However, what accounts for this enriched propensity is not known. Here we show that specific arrangements of polyA/T tracts, which are abundant and enriched in yeast intergenic regions explain this observation. We provide evidence that polyA/T tracts, which are known to act as Nucleosome Depleted Regions in a regulatory context, have been coopted through de novo gene emergence for the evolution of novel small genes encoding proteins with predicted transmembrane domains. Our findings contribute to our understanding of the process of de novo gene evolution and show how seemingly distinct but interacting levels of functionality can exist within the same genomic loci.

## Introduction

Entirely novel protein-coding genes can originate de novo from previously non-genic genomic regions^1,2^. This realization has raised the question of whether the properties and patterns found within such non-genic regions influence the evolutionary trajectories that de novo genes can follow. A recent study showed that intergenic regions of the budding yeast genome have a surprisingly high potential to, if translated, encode proteins with transmembrane (TM) domains^3^ as predicted by Phobius^4^. This observation is partly due to the high AT-content of such intergenic regions, which does correlate with hydrophobicity and thereby increases a peptide’s propensity to form a TM domain. However, the observed frequency of predicted-TM domains in intergenic regions still exceeds expectations for their given AT-content. Thus, intergenic regions are described to have an enriched propensity towards encoding TM domains. In addition, evolutionarily young genes in *S. cerevisiae* encode significantly more TM domains than expected given their AT-content, as well as more TM domains relative to conserved genes. TM-enrichment in intergenic regions is not a property restricted to *S. cerevisiae*, as we later showed that it is prevalent in the vast majority of Saccharomycotina yeast genomes^5^. Our previous work excluded a number of genomic properties, locations, and the presence of promoters based on neighboring gene orientation, as likely explanations of TM-enrichment. As that study stopped short of proposing a plausible explanation, we sought to discover specific motifs that might explain this cryptic property here.

## Results and Discussion

We asked whether there are any underlying DNA motifs within yeast intergenic regions that may influence the emergence of transmembrane (TM) domains. We obtained intergenic regions in which hypothetical in-frame stop codons have been removed (iORFs) and TM domains predicted by Phobius from Tassios et al. (N=6,384 total; iORFs with TM domains [TM+]: 3,544; iORFs without TM domains [TM-] 2,628; see **Methods**). Using Multiple Em for Motif Elicitation^6^ (MEME), we identified three enriched motifs in the iORFs consisting of 1) a tract of consecutive A’s (polyA), 2) a tract of consecutive T’s (polyT), and 3) a tract of AT-repeats (TATA) (**Figure 1A**) (see **Methods**). The polyA motif was present in 49.7% of iORFs, polyT in 47.5%, and TATA in 6.7%. While there is a weak enrichment of the polyA motif within TM+ iORFs (52.4% compared to 45.5% of TM-ones; Chi-squared P=0.00185), the polyT motif is found in 64% of TM+ iORFs but only 24% of TM-iORFs (Chi-squared P=1.066533e-83). The TATA motif is also enriched, found in 8.4% of TM+ vs. 4.7% of TM- (Chi-squared P=1.336e-07; **Figure 1B**). Enrichment of the polyT motif in TM+ iORFs is even stronger when considering all motif occurrences normalized by total length, as identified with Find Individual Motif Occurrences (FIMO; Supp. Figure 1A). While more frequent near the edges, all three motifs are found across the entire length of iORFs (Supp. Figure 1B) including when only considering iORFs longer than 500nt (Supp. Figure 1C). Note that when MEME analysis is performed separately on TM+ and TM-iORFs, the polyT motif is not even detected in TM-. Differential motif enrichment analysis further supports these results (see Methods and Supp. Figure 1D for motifs). To summarize, we demonstrate that intergenic regions predicted to encode TM domains are significantly enriched in polyA/T repeats.

**Figure 1.**
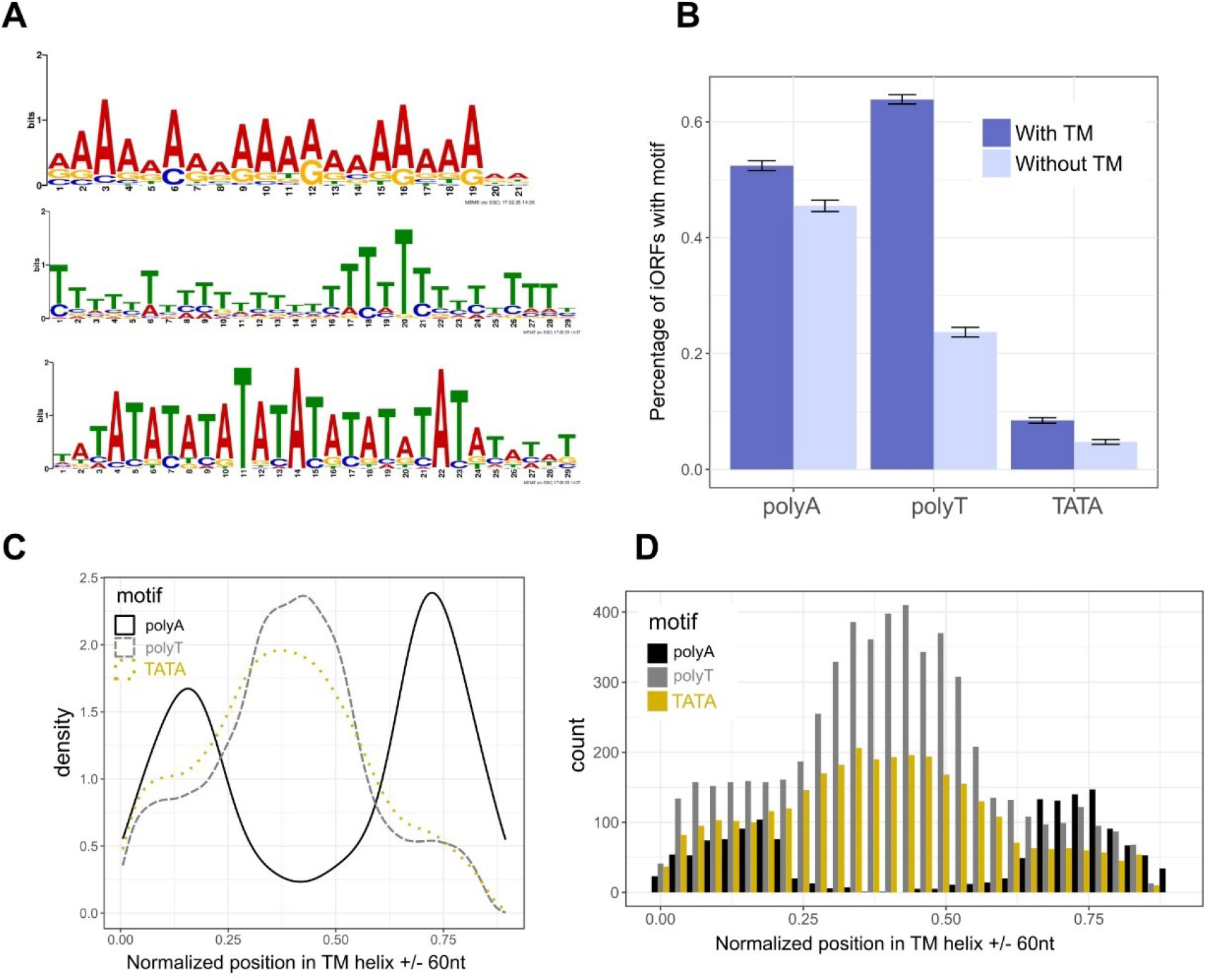
**A:** Significantly enriched DNA motifs in artificial iORFs. Top: E-value=3.1e-089, Sites=1,829, Width=29; Middle: E-value= 2.0e-069, Sites=2,503, Width=29; Bottom: E-value=2.4e-031, Sites=92, Width=38. (a fourth enriched motif only found in 6 sites is not shown) **B**: Percentages of iORFs with and without predicted TM domains, that have the polyA, polyT and TATA-containing motif seen in **A**, based on a FIMO motif search. **C**: Density distributions of the position of the three motifs (n=29,738) across each iORF TM-encoding segment +/-60nts, with 0 representing the start of the sequence and 1 its end. **D:** Same as C but only for sequences with at least one occurrence of at least two motifs (n=10,418) and distributions are shown as histograms.

To support the link between the presence of these motifs and predicted TM domains, we asked how many of the motifs occur specifically within the segments of TM+ iORFs which would encode a TM helix using Phobius, as well as the 60nt regions up and downstream of them (since the context of a predicted TM domain is taken into account by the prediction model). While predicted TM-helix encoding segments constitute only 34.6% of the total iORF length (937,918/2,711,967nt) they contain 78% of detected polyT motif nucleotides (637,420/813,711nt; two-proportion Z-test, 95% confidence, P<1e-16), 64% of TATA motifs (197,548/307,255nt; two-proportion Z-test, 95% confidence, P<1e-16), but only 31% of polyA (76,580/246,920nt; two-proportion Z-test, 95% confidence, P<1e-16). These proportions are similarly enriched when only considering motifs that overlap less than 50% (thus keeping only 10,361 out of the total 51,000 occurrences: 66%, 54.5%, 31%). A FIMO scan of only the TM helix-encoding segments +/-60nt found that out of the 6,606 TM-helix encoding segments, 3,815 (58%) have the polyT motif, 1,578 (28%) have the TATA motif, and 4,449 (67%) have at least one of the two. Finally, all three motifs can also be directly detected by MEME on the 6,606 sequences (Supp. Figure 1E). Therefore, polyT and TATA-containing motifs explain the elevated potential of iORFs to encode TM domains.

When examining the distribution of the motifs detected on the TM-encoding regions, we observed a clear pattern, in which the polyA motif flanks either the polyT or TATA-containing motif. **Figure 1C** shows this distribution of all three motifs along the TM-encoding sequences +/-60nt, and **Figure 1D** shows this distribution of motifs on sequences with at least two of the motifs. The yeast intergenic regions are known to be AT-rich^7^ specifically with AA/AT/TA/TT dinucleotide pairs^8^, and `TTTT`, `AAAA`, and `TATA` 4-mers^9^. However, to the best of our knowledge, the arrangement of such large tracts of polyA, polyT, and TATA motifs in this particular order has not been described in yeast intergenic regions. Yet it is this specific arrangement that underlies the potential of iORFs to encode TM domains. We further found that co-occurrence of polyT and polyA motifs is statistically significant in TM+ iORFs, with 77% iORFS being TM+ when motifs co-occur compared to 70% when only polyT occurs (Chi-squared P=2.08e-05).

Given our previous findings about trends in TM propensity (i.e. high in iORFs, lower in young genes, even lower in conserved genes) we hypothesized that since these motifs underlie TM enrichment, traces of them should also exist in de novo genes inferred to have originated from intergenic regions. We thus analyzed a set of 37 de novo genes which we previously identified using a robust ancestral sequence reconstruction pipeline^10^ focusing on the 28 that do not overlap other genes more than 10% of their length. Note that the proteins encoded by 43% (12/28) of these genes are predicted to have a TM domain. We searched with FIMO using the 3 motifs as queries and identified them in 16 (polyT), 3 (polyA) and 0 (TATA) sequences (see **Methods**). Out of the 12 de novo proteins with a TM domain, 9 (75%) have a match to the polyT motif and these matches fall within the TM-encoding segments +/-60nt. In other words, we find that de novo genes are also enriched with the polyT motif which likely predates their emergence.

To further strengthen the hypothesis that the motifs predated the origination of these genes we did a search using the polyT motif as query, within the orthologous regions of these de novo genes in the other 7 Saccharomyces species and were able to find them in at least one orthologous region of 21/28 de novo genes (88/258 orthologous sequences with a match overall; 642 matches in total). Therefore, this motif is even more prevalent within the orthologous non-coding regions than the de novo genes themselves. This finding is consistent with the orthologous non-coding regions more closely reflecting the ancestral non-coding status of the sequence, as the gene likely diverged through the accumulation of non-neutral mutations. Furthermore, the motif is also present in other sets of previously published putative de novo *S. cerevisiae* – specific genes, found by FIMO in 50% (87/172) sequences (see **Methods**). Importantly, no similar motif could be detected by MEME in *S. cerevisiae* genes that are at least as ancient as Fungi (n=4,477) and FIMO only finds it in 6 genes (0.13%, Chi-squared P<1e-16 when comparing to the proportion in the 28 de novo genes). Thus, this motif is specifically associated with candidate de novo *S. cerevisiae* genes.

The presence of polyA/T tracts within intergenic regions in the yeast genome has been known since the 1980s and there is a body of evidence supporting their role as Nucleosome Depleted Regions (NDRs) which function near promoters to enable and regulate gene expression^7,11,12^. Assuming that the original function of these motifs, that is what they originally became selected for during evolution, is to act as NDRs, it seems that they were primed to assume an entirely new function within emerging protein-coding sequences. Interestingly however, as we previously showed^5^, TM enrichment, and also polyA/T motifs (see Supp. Figure 1B and C), are not limited to promoter regions but are also found close to the center of intergenic regions, and we would not expect this if these motifs were strictly promoter associated. To our knowledge, this is the first reported occurrence of a specific presumably regulatory motif directly influencing the evolution of novel genes. At the same time, our findings provide additional support for the de novo origination of 28 previously described de novo genes (and potentially more across the other datasets), as the polyA/T motif can be viewed as an abundant intergenic signature that is unlikely to have other origins.

## Materials and Methods

### Sequence datasets

The artificial intergenic ORFs (iORFs) used in this analysis were generated by ^5^ and are the same used in ^3^ They consist of entire intergenic regions of the *S. cerevisiae* reference genome and annotation, out of which in-frame stop codons have been removed (only the +1 frame is considered). We started with 6,384 such iORFs over 50nt length. Based on TM-prediction by Phobius^4^, we split the dataset into 3,544 iORFS with at least one TM domain (TM+) and 2,628 iORFs without TM domains or signal peptides (TM-; 212 iORFs with only signal peptides were removed to allow a cleaner comparison). We parsed the Phobius output to extract the coordinates of the sequence that encodes each TM helix and we extended that region up and downstream 60 nucleotides or until the start/end of the next/previous TM helix.

The set of 37 robust de novo gene candidates was taken from ^10^ and contains those genes for which no version of ancestral sequence reconstruction resulted in the presence of a statistically significant ancestral ORF. These were inspected on the genome browser of Saccharomyces Genome Database^13^ for overlap to conserved genes on the other strand. Additional sets of *S. cerevisiae*-specific putative de novo genes come from Carvunis et al.^14^, Vakirlis et al.^15^ and Wu and Knudson^17^ (the dataset of Blevins et al.^16^ was not included as it mostly contains overlapping genes). We kept genes that were reported to be species-specific by at least one of the three studies and were not predicted to be older by any of the three. This resulted in 189 non-redundant genes. To further confirm their species-specific status we conducted a similarity search against all fungal RefSeq proteins using BLASTP and we removed 17 proteins with a match with E-value<0.01 in a non-*S. cerevisiae* protein. *S. cerevisiae* reference annotation (R64) protein-coding genes with origins at least as ancient as Fungi (n=4,477) were predicted using broad sequence similarity searches and phylostratigraphy as in ^19^.

### Motif analyses

Enriched motifs were identified using a local installation of MEME with the following command: *meme [INPUT_FILE] -dna -oc*. *-nostatus -time 14400 -mod zoops -nmotifs 5 -minw 6 -maxw 50 -objfun classic -markov_order 0 -p 10*

Differential motif analysis was performed with the following command:

*meme [iORFs_WITH_TM_DOMAINS] -dna -oc*. *-nostatus -time 14400 -mod zoops -nmotifs 3 -minw 6 - maxw 50 -objfun de -neg [iORFs_WITHOUT_TM_DOMAINS]-markov_order 0*

Motif scanning was performed with FIMO with the following command:

*fimo --oc*. *--verbosity 1 --bgfile --nrdb thresh 1*.*0E-4 --norc [MOTIF] [SEQUENCES_TO_SCAN]*

Only motifs with a q-value < 0.01 were considered statistically significant. All motif analyses were restricted on the given (forward) strand of the sequences.

## Acknowledgements

The research project was supported by the Hellenic Foundation for Research and Innovation (H.F.R.I.) under the “3rd Call for H.F.R.I. Research Projects to support Post-Doctoral Researchers” to N.V. (Project Number:7330) and by a G4 grant from the Pasteur Institute awarded to N.V. T.F. is supported by a University of Zurich Postdoc Grant (FK-23-120).

---

**Supplementary Figure 1.**
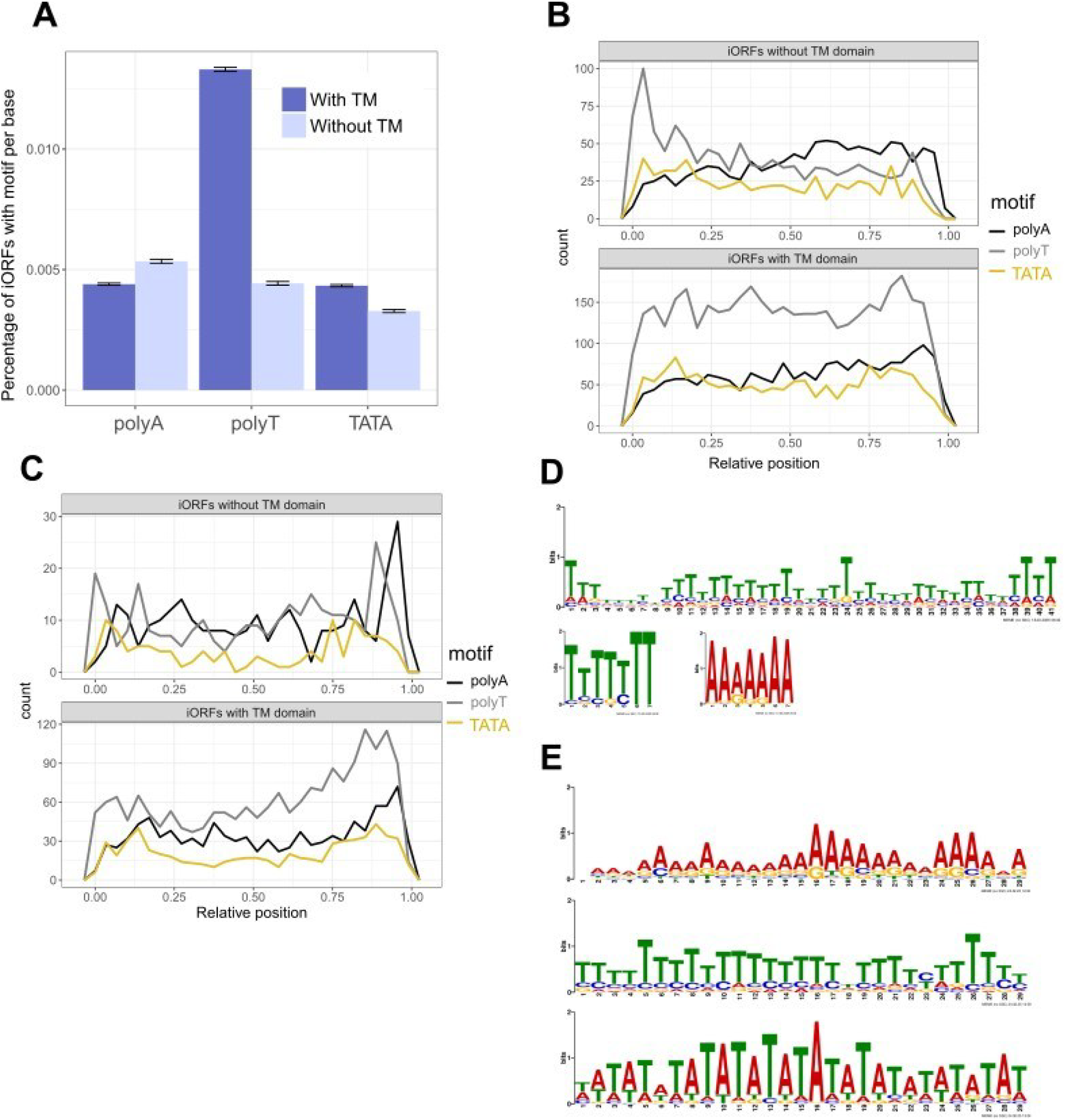
**A:** Percentage of motif occurrences of the polyA (N=15,210), polyT (N=28,059) and TATA-containing motif (N=12,270) per base (that is, normalized by total sequence length), on iORFs with (total length 1,844,901nt) and without predicted TM domains (total length 793,350). **B**: Distribution of normalized locations of motifs along 3,544 iORFs with TM domains and 2,628 iORFs without TM domains. **C**: Same as B but only on iORFs longer than 500nt (top n=346; bottom n=1,161). **D**: Top three differentially enriched DNA motifs in artificial iORFs with predicted TM domains compared to those iORFs without predicted TM domains. Top: E-value= 3.4e-300, Sites=2660, Width=41; Bottom left: E-value=1.4e-042, Sites=2949, Width=7; Bottom right: E-value=1.3e-020, Sites=2884, Width=7. **E:** Top three significantly enriched DNA motifs in TM helix encoding regions +/-60nt, of artificial iORFs with predicted TM domains. Top: E-value=2.5e-181, Sites=1,113, Width=29; Middle: E-value= 1.2e-100, Sites=1,485, Width=29; Bottom: E-value=3.5e-062, Sites=508, Width=29.

